# Targeted memory reactivation during sleep to strengthen memory for arbitrary pairings

**DOI:** 10.1101/447623

**Authors:** Iliana M. Vargas, Eitan Schechtman, Ken A. Paller

**Affiliations:** Department of Psychology, Northwestern University, Evanston, IL 60208, USA

**Author notes:** Corresponding author, Department of Psychology, Northwestern University, Evanston, IL 60208, USA.

**Keywords:** consolidation, slow-wave sleep, learning, spatial memory

## Abstract

A powerful way to investigate memory consolidation during sleep utilizes acoustic stimulation to reactivate memories. In multiple studies, Targeted Memory Reactivation (TMR) using sounds associated with prior learning improved later memory, as in recalling locations where objects previously appeared. In the present experiment, we examined whether a variant of the same technique could strengthen memory for the locations of pairs of objects. Each sound was naturally connected to one object from each pair, but we hypothesized that both memories could be improved with TMR. We first asked participants to memorize each of 50 pairs of objects by associating the two objects with each other and with the sound of one of the objects (e.g., cat-meow). Next, objects were presented in unique locations on a grid. Participants learned these locations in an adaptive procedure. During an afternoon nap, 25 of the sounds were quietly presented. In memory tests given twice before and twice after the nap, participants heard the sound for each object pair and were asked to recall the name of the second object and the locations of both objects. Forgetting scores were calculated using the mean difference between pre-nap and post-nap spatial recall errors. We found less forgetting after the nap for cued compared to non-cued objects. Additionally, the extent of forgetting tended to be similar for the two members of each pair, but only for cued pairs. Results thus substantiate the potential for sounds to reactivate spatial memories during sleep and thereby improve subsequent recall performance, even for multiple objects associated with a single sound and when participants must learn a novel sound-object association.

**Highlights:**

- Memories can be improved during sleep using arbitrary sounds
- Participants learned a random screen location for each of 100 objects
- Objects were learned in pairs with the characteristic sound of one of the objects
- Half of those sounds were presented during a nap that followed learning
- After sleep, location recall was more accurate for cued than for non-cued objects

## Introduction

Sleep is widely thought to be important for memory consolidation. A contemporary theoretical framework for work in this area is based on the idea that, during sleep, memories are reactivated while particular patterns of neural activity are recapitulated or replayed (Rasch & Born, 2013). This replay and associated plasticity in hippocampal-neocortical networks may be essential for memory consolidation. Yet, much remains to be elucidated about these consolidation mechanisms.

The first neural evidence for memory reactivation during sleep came from rodent studies in which place cells active during learning showed the same temporal order of activation during subsequent sleep (Pavlides & Winson, 1989; Wilson & McNaughton, 1994). However, these studies did not show whether subsequent memory performance changed as a function of place-cell reactivation during sleep. In humans, the strongest evidence for memory reactivation during sleep comes from a procedure called Targeted Memory Reactivation (TMR). With this procedure, investigators choose which memories to reactivate and then monitor the influence of reactivation on subsequent memory performance (Oudiette & Paller, 2013; Schreiner & Rasch, 2015). Presumably, cues presented during sleep engage replay as well as modification of memories associated with a previously learned task. Initial research on strengthening object-location memory with TMR provided evidence of reactivating an entire learning session using odors (Rasch et al., 2007) and of reactivating specific object-location associations using sounds (Rudoy et al., 2009). Although TMR is effective at strengthening memory in a variety of different tasks beyond object locations (Schouten, Pereira, Tops, & Louzada, 2017), the extent to which single auditory cues can be used to reactivate spatial memories encompassing more than one spatial association, and going beyond pre-existing sound-object associations (e.g., meow-cat), remains unexplored.

Here, we asked whether TMR could be used to reactivate and strengthen memory for locations of pairs of objects associated with a single sound. We hypothesized that multiple object-locations could be reactivated at the same time. Reactivating complex associations is a first step in exploring the selective reactivation of multiple memory items using a single sound. The results provide information about the future potential of sleep reactivation. For instance, successful reactivation of unrelated pairs of objects in this experiment would open the door to more elaborate strategies to reactivate a larger number of distinct memories with an individual sound.

## Materials and Methods

Participants were members of the Northwestern community (*N* = 24, ages 18-24 years) with no known history of neurological disease who claimed to be able to nap in the afternoon. Participants were instructed to wake up 2 hours earlier than usual and not have any caffeine the day of the experiment. Results do not include data from an additional 25 participants (20 failed to enter NREM sleep long enough for one round of cue presentation; 3 reported hearing the sounds during the nap; 1 dropped out of the study before the nap; and 1 was excluded due to below 50% accuracy on the cued recall test). The Northwestern University Institutional Review Board approved the procedure.

The experiment consisted of six phases, as shown in Figure 1: (1) learning pairs, (2) learning locations, (3) practicing pairs and locations, (4) pre-nap test for pairs and their locations, (5) 90-min nap opportunity, and (6) post-nap test for pairs and their locations. The electroencephalogram (EEG) was recorded during the testing and nap phases of the experiment. EEG was recorded from 21 scalp locations from the 10-20 system (Fpz, Fp1, Fp2, Cz, C3, C4, F3, F4, F7, F8, Pz, P3, P4, T3, T4, T5, T6, Oz, O1, O2) and both mastoids. Additional electrodes were placed on the face for recording vertical and horizontal electro-oculogram (EOG) and chin electromyogram (EMG). Electrodes were referenced to the left mastoid electrode and rereferenced offline to the average of the two mastoids. Impedances were brought down to 5kΩ and voltage was sampled at 1000 Hz.

**Figure 1.**
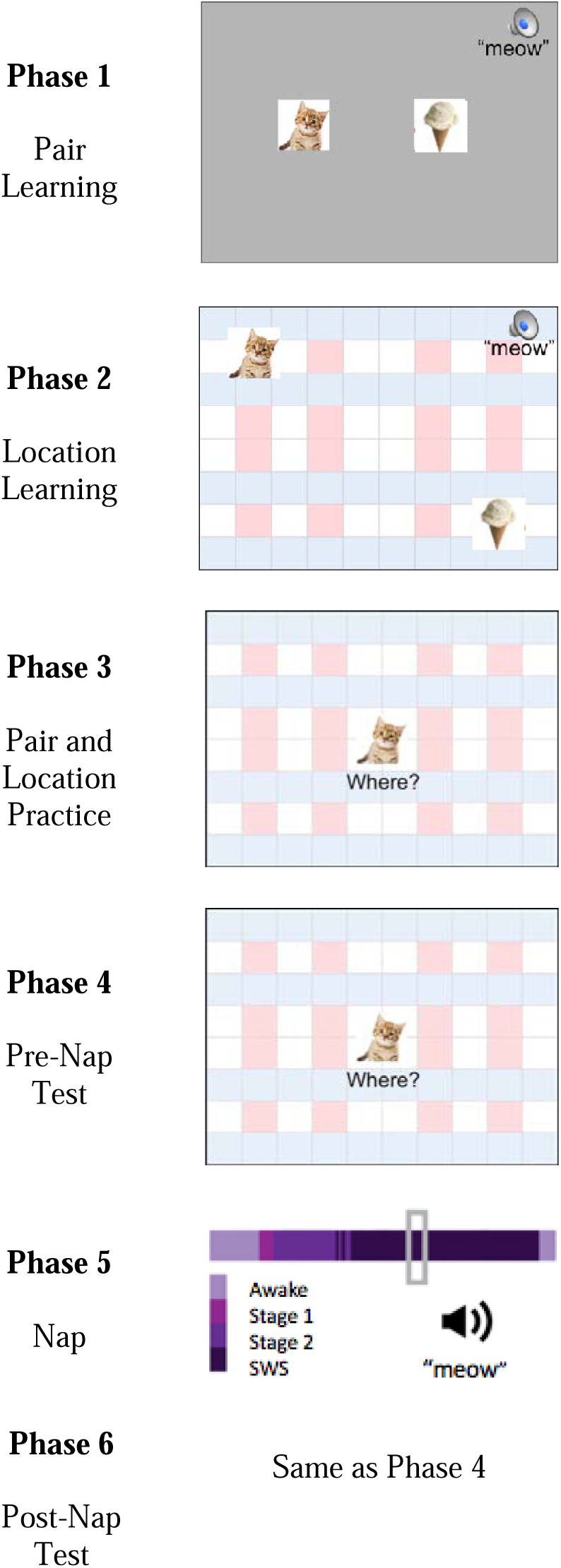
Schematic depiction of experimental timeline.

Stimuli consisted of photographic images of 100 common objects (Table 1), half with a characteristic sound lasting less than 500 ms and presented through a speaker. Images were 150 × 150 pixels (5.3 cm × 5.3 cm) and presented on a grid background screen of 1,000 × 800 pixel (35.7 × 28.6 cm) from a distance of 100 cm. Each of 50 objects with a sound (Object A) was randomly paired with one of 50 objects without a sound (Object B). Pair combinations were randomized for each participant.

**Table l.**
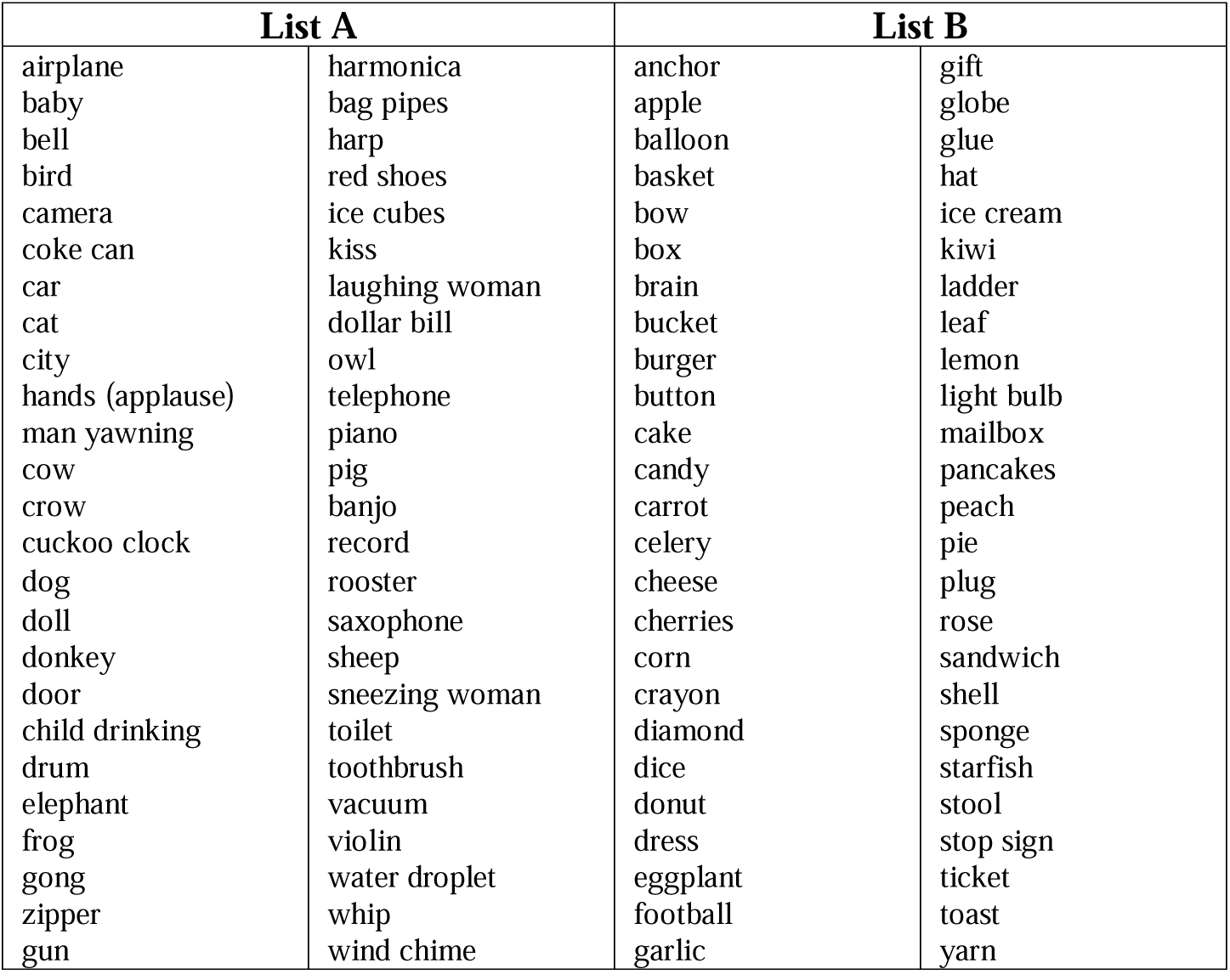
Lists of stimuli used as Objects A and B

In Phase 1, participants were instructed to memorize object pairs. Each of 50 A-B pairs appeared on a gray background on the screen (Object A to the left of Object B) for 5000 ms, followed by a 500-ms interstimulus interval. Participants were advised to construct a story for how the two objects might be related. They were also told to pay close attention to each sound, which was presented along with the stimuli, and to associate it with both objects.

In Phase 2, participants were instructed to memorize locations of objects presented on a grid background. Two objects were simultaneously presented in different locations on the screen. Each pair appeared on the screen for 6000 ms, accompanied by a single presentation of the associated sound. This was followed by a 500-ms interstimulus interval. The location for the center of each object was assigned using an X-Y coordinate system. A coordinate of (0, 0) corresponded to the center of the screen. The X and Y coordinates for Object A were randomly assigned values between −300 to 300. For Object B, one coordinate (either X or Y) was assigned a random value between −300 to 300. If Object A was within 210 pixels from the boundary of the 600 × 600 pixel area in which stimuli could potentially appear, then the second coordinate for Object B was chosen at a random location between Object A and the opposite boundary; otherwise the object was assigned a random location on one side of Object A or the other, within the range of −300 to 300. In all cases, Objects A and B were at least 210 pixels from each other so as not to overlap.

In Phase 3, participants were asked to place objects in their correct locations and to recall Object B when prompted with Object A. Repeated location practice was conducted using a drop-out method. In each trial, Object A appeared in the middle of the screen for 1000 ms along with the word “Where?” The associated sound was presented at the same time as the object. After 1000 ms, “Where?” disappeared from the screen and participants indicated their choice using a mouse by dragging the object to a location and making a left click. The object then disappeared for a 100-ms interval and appeared in its correct location for 3000 ms, again accompanied by the associated sound. A prompt asking “What was its pair?” appeared for 1000 ms while Object A was still displayed on the screen. Participants used a keyboard to type out the name of Object B. Once they entered the name or indicated that they did not know the pair by typing “idk,” the sound was presented again and Object B appeared in the middle of the screen for 1000 ms along with the word “Where?” After the word disappeared, participants indicated their choice by dragging the object to a location and making a left click. The object disappeared for 100 ms and then reappeared in its correct location for 3000 ms, accompanied by the sound. Object A remained on the screen through the duration of each trial. For the purpose of practice in this phase, a placement within 150 pixels of the correct location was considered a correct response. In each run through the list, the pairs were tested in the same order. All 50 pairs appeared in the first two runs. Thereafter, pairs were dropped out as follows. If both Object A and Object B were placed in the correct location twice, that pair did not appear again. Otherwise, the pair was included in the subsequent run. The location did not have to be correct on consecutive runs for the object to be dropped. If neither object location was correct, the pair was included in the same way. If only the location for Object A was correct twice, Object A appeared in its correct location and the trial continued from there (prompt for name of Object B and then practice for location of Object B). If only Object B was correct twice, the trial transpired as usual except that Object B appeared in the correct location rather than requiring location recall for Object B. Participants were allowed to take a short break between runs if needed. After all objects were placed in the correct location twice, Phase 3 ended.

In Phase 4, there was a pre-nap test for the 50 paired object names and 100 object locations. There were two runs with a different random order each time. The testing format was similar to the practice except that there was no feedback. Object A was presented in the middle of the screen for 1000 ms with the word “Where?” and the associated sound. Once the word “Where?” disappeared, participants attempted to drag the object to its correct location. Next, they saw the question: “What was its pair?” for 1000 ms. After they typed in the name, Object B appeared in the middle of the screen for 1000 ms with the word “Where?” and the associated sound. Participants attempted to drag the object to its correct location. Object A remained on the screen for the duration of each trial. Each trial was followed by a 500-ms interval when only the grid was displayed.

Phase 5 included a 90-minute nap opportunity that began approximately 2 hours after the beginning of the study. The nap took place in the same room as earlier phases. The futon chair used in the prior phases was converted into a bed, with sheets and a pillow. Participants reclined while listening to white noise. Speakers were placed on a shelf to the left and right of their head and sound intensity was approximately 45 dB for the white noise and 50 dB for the individual sounds. After lights were turned off, the participant attempted to sleep. Sleep stages during the nap were monitored online using continuous EEG, EMG, and EOG recordings. When the experimenter determined that SWS had been reached, or that it might not be reached and that presenting sounds during Stage 2 would not disturb sleep, half of the sounds from learning were presented repeatedly. Sounds were presented in a random order in each presentation of the list. These specific 25 sounds were selected by first taking the distance between the recalled location and the correct location, averaged across both objects and both test runs (see Phase 4). Pairs were ranked based on performance and either the even- or odd-ranked sounds were played during sleep. Stimulation rate was one sound every 5800 ms and the sounds continued until slow-wave sleep ended. For most participants, cues were presented during both SWS and Stage 2 (*n*=16). Two participants did not reach SWS and were only cued during Stage 2, and six participants were only cued during SWS. Each sound was presented during sleep 1-19 times (mean = 7). Participants were allowed to wake up naturally or were woken up after 90 min. Then, a 10- to 15-minute break ensued.

In Phase 6, participants were tested on the object names and locations in the same manner as in the pre-nap test. Prior to the test, they filled out the Karolinska Sleep Log, which assesses the quality and length of the previous night of sleep. After the post-nap test, they completed a questionnaire to assess the difficulty of the task, nap quality, and expectations about the experiment.

### Sleep Physiology

After the experiment, continuous EEG was down-sampled to 128 Hz and filtered at .5-50 Hz using an infinite impulse response Butterworth filter. Sleep stages were formally identified using standard sleep scoring criteria (Iber et al., 2007).

Standard analyses of sleep oscillations were computed focusing on two clusters of interest (frontal using Fpz, Fp1, Fp2; central-parietal using Cz, Pz, C3, C4, P3, P4). A fast Fourier transform using a Hanning function and 5 second intervals was performed on NREM sleep epochs. We extracted mean power for delta (1- 4 Hz) and sigma (12 −15 Hz) bands. For slow-oscillation analyses, EEG was low-pass filtered at 3.5 Hz. Slow oscillations were detected by finding adjacent points in which the EEG signal voltage changed from positive to negative that were .5-2.0 s apart from each other, and when the maximum peak-to-peak amplitude between the two points was greater than 75 μV. Spindles were automatically detected (Mölle et al., 2011) by first filtering EEG data between 11-16 Hz and calculating root mean squared (RMS) voltage using a sliding 200-ms window. A spindle was counted if the RMS crossed a threshold of 1.5 standard deviations of the signal and remained above the threshold for 0.5-3.0 s. Because fast spindles and slow spindles show different topographies, with fast spindles predominant at parietal and central locations and slow spindles at frontal locations, probably with distinct neural generators (Rasch & Born, 2013), we separately analyzed fast (> 13.5 Hz) and slow spindles (< 13.5 Hz).

### Behavioral Data

Behavioral data for pairs in which the participant was unable to recall the name of Object B on the first post-nap test were excluded from analysis. We also excluded trials with poor pre-sleep spatial learning if both the average pretest error was more than 212 pixels and the two recalled pretest locations were more than 212 pixels away from each other (212 pixels is the length of the picture’s diagonal). That is, if the object was not placed close to the original location and also was placed inconsistently, then the location was presumably not effectively learned. After the exclusion of these trials, the average number of trials per participant was 97 ± 1 (mean ± *SD*, maximum 100). Because the correlation analysis requires data for both A and B objects, for this analysis the average number of pairs per participant was 48 ± 1. For spatial recall data presented below, the standard error of the mean across participants was computed after averaging scores across both pre-nap or both post-nap tests for each individual.

The main analysis concerned the change in recall error for objects cued by a sound during sleep compared to objects that were not cued during sleep. In particular, we hypothesized that TMR would reduce forgetting for both objects in a pair if the associated sound was presented during sleep. Recall error was computed as the log-transformed distance to the studied location, averaged across the two pre-nap or two post-nap tests. A forgetting score for each object was calculated as the average error at post-nap test minus average error at pre-nap test. A higher score indicates more forgetting after the nap. To test statistical significance, forgetting scores were submitted to an ANOVA with trial type (A/B) and cuing (cued/not cued) as within-subject factors. A cuing advantage score was calculated for each participant as the difference in forgetting score for not-cued objects minus cued objects (higher score indicates larger relative benefit due to cuing). The cuing advantage score was used to investigate the relationship between behavioral measures and sleep physiology.

In addition to forgetting scores, we also evaluated the within-test consistency with which object locations were recalled, as another measure of the quality of learning (better learning should produce more consistent location recall responses). This consistency index was calculated as the (log transformed) distance between the first and second placement of each object on the two runs of the pre-nap test, or on the two runs of the post-nap test. A lower number indicates greater recall consistency. To test statistical significance, the within-test consistency scores were submitted to an ANOVA with time (pre-nap test/post-nap test), object type (A/B), and cuing (cued/not cued) as within-subject factors.

We were also interested in location recall consistency within pairs. We hypothesized that TMR might conjointly improve memory for both objects in a pair, such that A and B objects in cued pairs would show similar changes in error. To calculate how error changed in paired trials, for each participant, we computed the correlation between error for A trials and error for B trials, before and after the nap, as well as the correlation between forgetting scores for A and B paired trials. Correlation scores were transformed to *z*-scores using a Fisher transformation to conduct hypothesis testing. A change in correlation due to cuing was obtained as the difference in change in correlation (*z*-transformed) between cued and not-cued pairs.

## Results

The training procedures in Phase 3 were effective, as participants were almost always able to recall the name of the second object. A perfect cued recall score was achieved by 17 participants. The average number of words missed was 1 ± 2 (mean ± *SD*) out of 50 in the post-nap test.

For spatial recall, the mean pre-nap error across both tests was 82.5 ± 3.0 pixels (30 mm) from the original location (Table 2). For reference, the length of each picture’s diagonal was 212 pixels (77 mm), so with this average magnitude of error the object would largely overlap with a perfect recall placement. Of course, some objects were recalled with less spatial accuracy, and some greater spatial accuracy, but none with absolutely perfect accuracy. When tested approximately 2 hours later, recall was still quite accurate. After the nap, objects were placed 87.3 ± 3.3 pixels (32 mm) from the original location.

**Table 2.** Behavioral data in the memory tests. 1 pixel = .367 mm

|  | **Error in Pixels**<br>**(All Trials)** | **A trials** | **B trials** |
| --- | --- | --- | --- |
| Average Pre-nap test Error | 82.5 ± 3.0 | 75.3 ± 2.3 | 89.9 ± 4.2 |
| Cued During Sleep | 84.9 ± 2.8 | 75.6 ± 2.6 | 94.3 ± 4.8 |
| Not Cued During Sleep | 80.9 ± 3.4 | 74.8 ± 2.3 | 85.4 ± 3.8 |
| Average Post-nap test Error | 87.3 ± 3.3 | 79.8 ± 2.6 | 94.9 ± 4.5 |
| Cued During Sleep | 87.9 ± 3.3 | 79.2 ± 2.7 | 96.8 ± 4.8 |
| Not Cued During Sleep | 86.7 ± 3.4 | 80.3 ± 3.0 | 92.9 ± 4.6 |
| Forgetting Score | 4.8 ± 0.9 | 4.6 ± 1.1 | 5.1 ± 1.1 |
| Cued During Sleep | 3.1 ± 0.9 | 3.6 ± 1.2 | 2.6 ± 1.2 |
| Not Cued During Sleep | 6.6 ± 1.4 | 5.5 ± 1.6 | 7.6 ± 1.7 |
|  | **Consistency**<br>**(in Pixels)** | **A trials** | **B trials** |
| Average Pre-nap test Consistency | 50.9 ± 2.7 | 48.8 ± 2.9 | 52.9 ± 2.8 |
| Cued During Sleep | 51.5 ± 3.0 | 50.4 ± 3.5 | 52.6 ± 3.2 |
| Not Cued During Sleep | 50.2 ± 2.9 | 47.1 ± 2.9 | 53.2 ± 3.1 |
| Average Post-nap test Consistency | 46.3 ± 3.0 | 44.1 ± 3.1 | 48.5 ± 3.1 |
| Cued During Sleep | 46.9 ± 3.1 | 44.4 ± 2.9 | 49.6 ± 3.7 |
| Not Cued During Sleep | 45.9 ± 3.4 | 43.8 ± 4.5 | 47.8 ± 3.1 |
| Change in Consistency | -4.5 ± 1.3 | -4.7 ± 1.9 | -4.4 ± 1.7 |
| Cued During Sleep | -4.6 ± 2.3 | -6.0 ± 2.6 | -3.2 ± 2.7 |
| Not Cued During Sleep | -4.3 ± 2.2 | -3.3 ± 3.2 | -5.4 ± 2.2 |
|  | **Correlation** |  |  |
| Correlation between A and B Pre-nap test Error | .19 ± .03 |  |  |
| Cued During Sleep | .22 ± .05 |  |  |
| Not Cued During Sleep | .20 ± .04 |  |  |
| Correlation between A and B Post-nap test Error | .16 ± .03 |  |  |
| Cued During Sleep | .24 ± .05 |  |  |
| Not Cued During Sleep | .09 ± .04 |  |  |
| Change in correlation between A and B Error | -.03 ± .04 |  |  |
| Cued During Sleep | .02 ± .06 |  |  |
| Not Cued During Sleep | -.11 ± .03 |  |  |
| Correlation between A and B Forgetting Scores | .13 ± .04 |  |  |
| Cued During Sleep | .19 ± .05 |  |  |
| Not Cued During Sleep | .05 ± .05 |  |  |

The chief hypothesis in this experiment was that memory would differ as a function of TMR during sleep. As shown in Figure 2, there was less forgetting for objects cued during sleep compared to objects that were not cued [*F*(1, 23) = 4.46, *p* < .05]. This cueing advantage was not significantly different between A trials and B trials [2.0 ±1.9 pixels and 5.1 ± 1.9 pixels, respectively; *F*(1, 23) = .75, *p* = .39]. There were negligible differences in forgetting between A and B trials, when collapsed across cuing conditions [4.6 ± 1.1 pixels and 5.1 ± 1.1 pixels, respectively, *F*(1, 23) = .47, *p* = .49].

**Figure 2.**
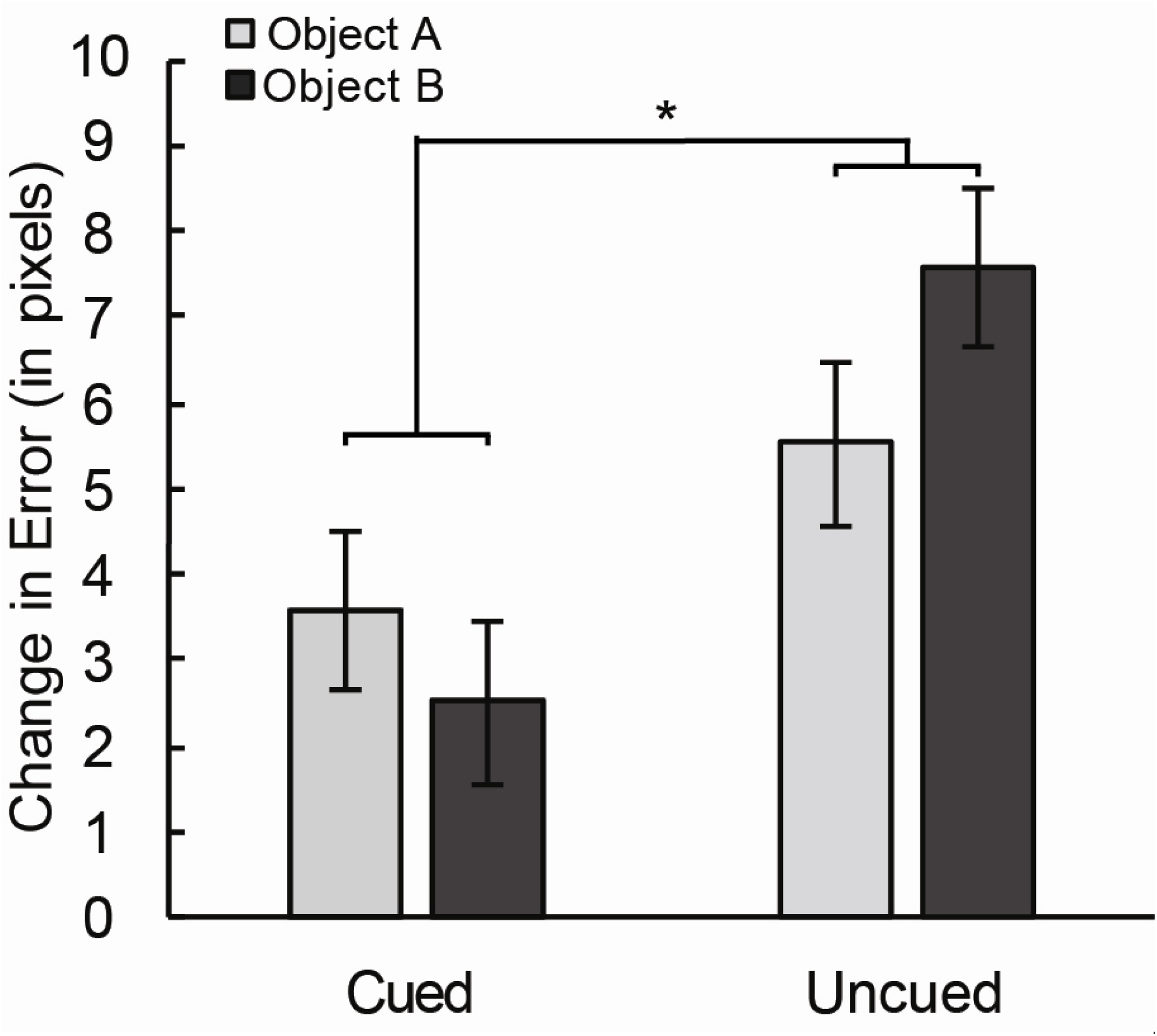
There was less forgetting from the pre-nap test to the post-nap test for object locations cued during sleep than for those not cued. This cuing benefit was not significantly different between Objects A and B. Error bars reflect ±1 standard error of the mean (SEM) for each condition adjusted to represent the variance for statistically evaluating the cuing benefit (i.e., SEM was calculated using each participant’s value for each condition after subtracting that participant’s mean value across all conditions). * − *p* < .05.

Within-test consistency, calculated as the distance between recalled locations for first and second placement of the same object on the same test, was better after the nap compared to before the nap, as shown in Figure 3 [*F*(1, 23) = 21.79, *p* < .001]. Participants were also more consistent in their placement of objects in A trials compared to B trials [46.5 ± 2.9 pixels vs. 50.7 ± 2.8 pixels, respectively; *F*(1, 23) = 26.80, *p* < .001]. However, TMR during sleep did not influence within-test consistency after the nap [*F*(1, 23) = .13, *p* = .72] or produce any changes in consistency as a function of A/B trial type [*F*(1, 23) = .37, *p* = .55]. Similar results were obtained when including the initial absolute error as a covariate, except that the difference between trials A and B was no longer significant. This additional analysis thus substantiated the notion that recall consistency within a test was greater after the nap.

**Figure 3.**
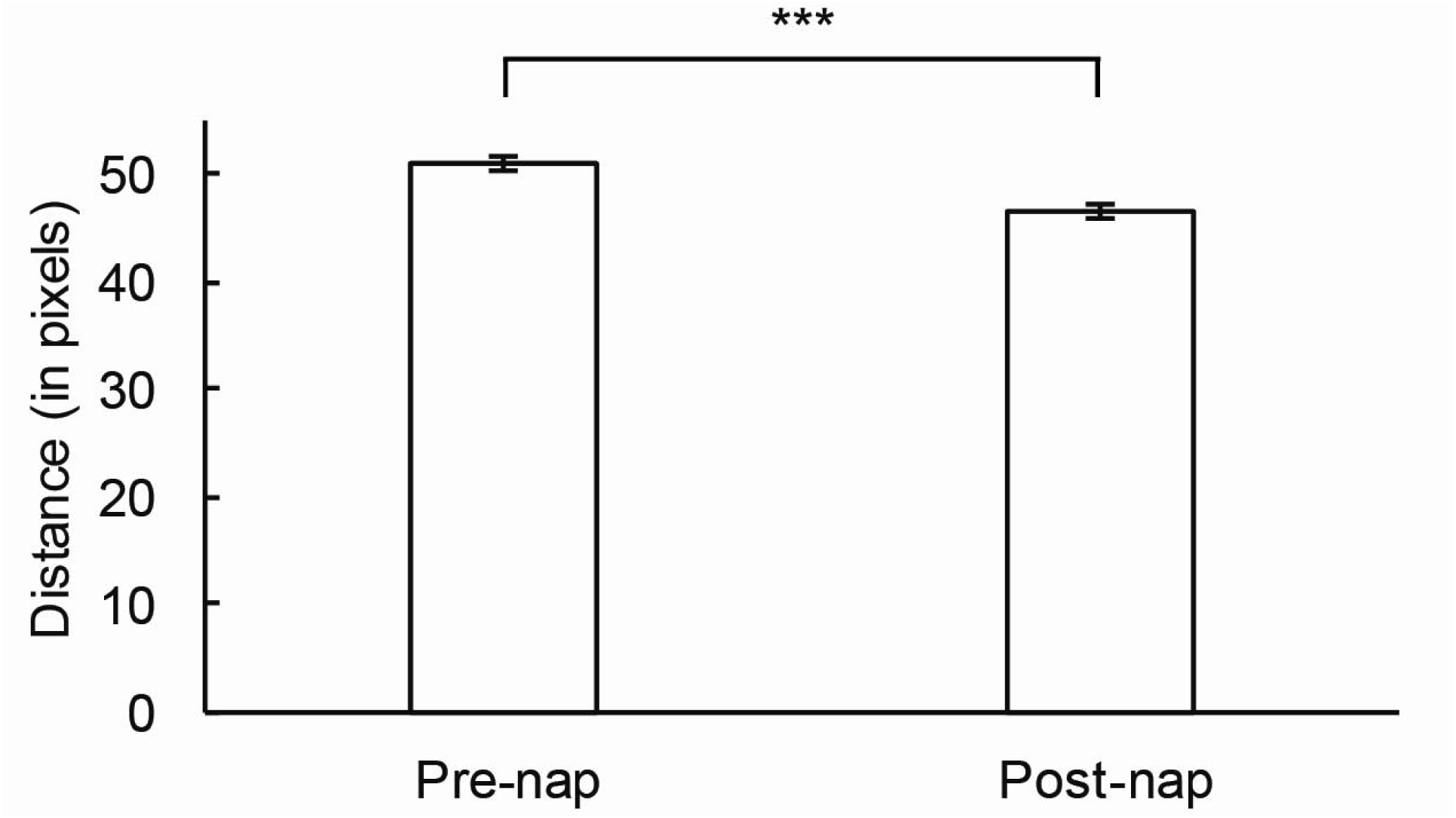
Participants attempted to recall each object location twice before the nap and twice after the nap, and recall consistency was measured as the distance between the two placements. The two placements were closer together after the nap compared to before the nap. Error bars represent ± 1 SEM (computed as in Figure 2). *** − *p* < .001.

Recall consistency can also be considered with respect to the relationship between errors for the A and B objects within each pair, given that both locations might tend to be forgotten or not forgotten together. The correlation between A and B error before the nap was .19 ± .03 (computed in each subject and averaged across subjects). As shown in Figure 4, the extent to which correlations changed from before to after the nap varied as a function of TMR [*F*(1,23) = 4.71, *p* < .05]. Specifically, pairs that were not cued showed a decrease in correlation after the nap [*t*(23) = 3.20, *p* = .004], whereas the correlation did not significantly change after the nap for cued pairs [*t*(23) = .47, *p* =.64]. That is, cuing enhanced the degree to which similar errors were made for A and B trials after sleep, and this similarity may indicate that A and B objects were reactivated together and benefitted in a correlated manner.

**Figure 4.**
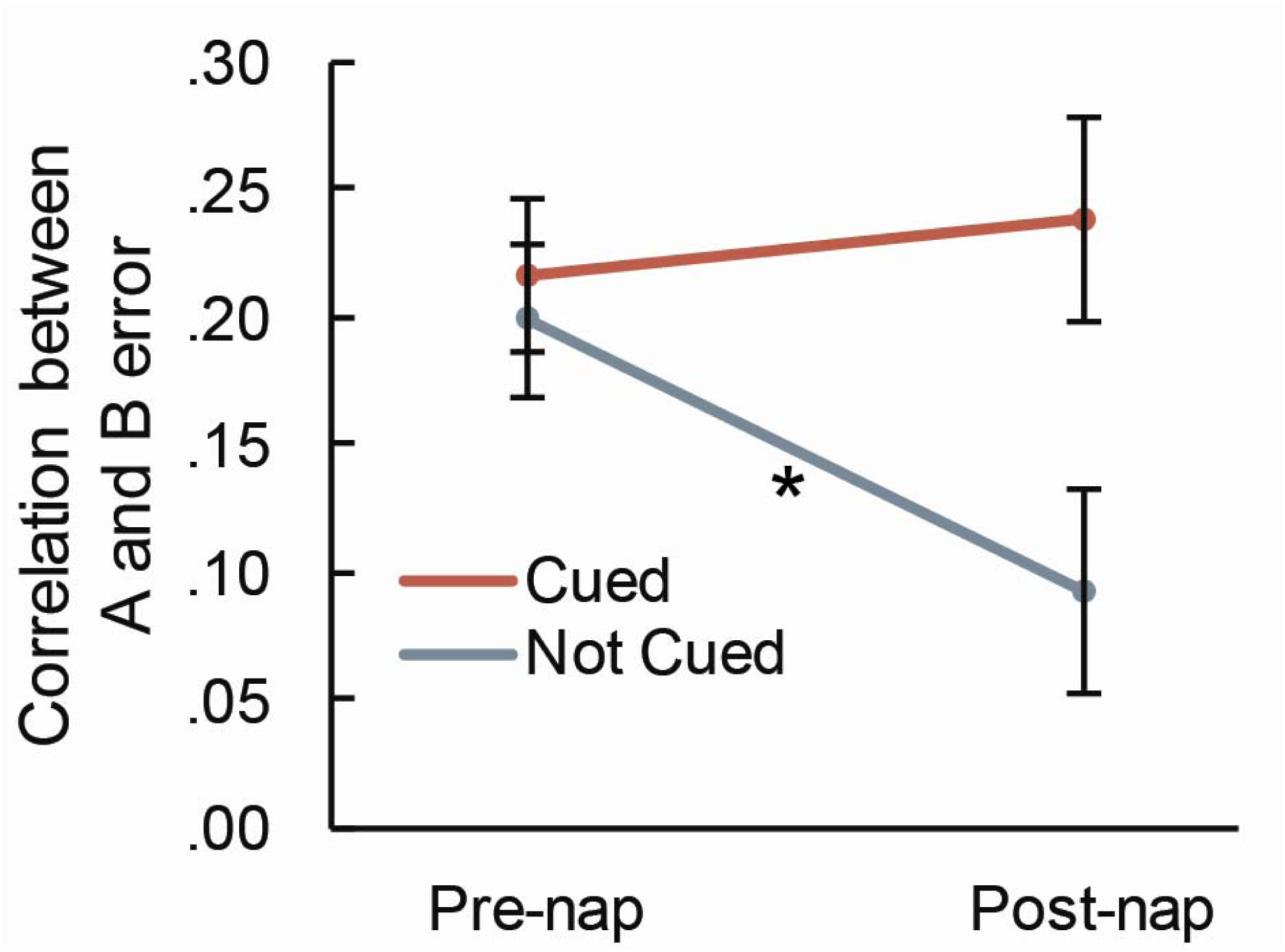
The correlation between A and B error significantly decreased after the nap for pairs that were not cued, but not for pairs that were cued. This pattern indicates that when a pair was not cued, the error changed at different rates for each object in a pair. Error bars represent ± 1 SEM (computed as in Figure 2). * − *p* < .05.

To further test this explanation, we considered the similarity between the level of forgetting over sleep for associated items and its dependence on cuing (Figure 5). There was a marginally greater correlation between A and B trial forgetting scores for cued versus not cued pairs [*t*(23) = 1.98, *p* = .06], suggesting that associated items shared similar forgetting patterns when cued.

**Figure 5.**
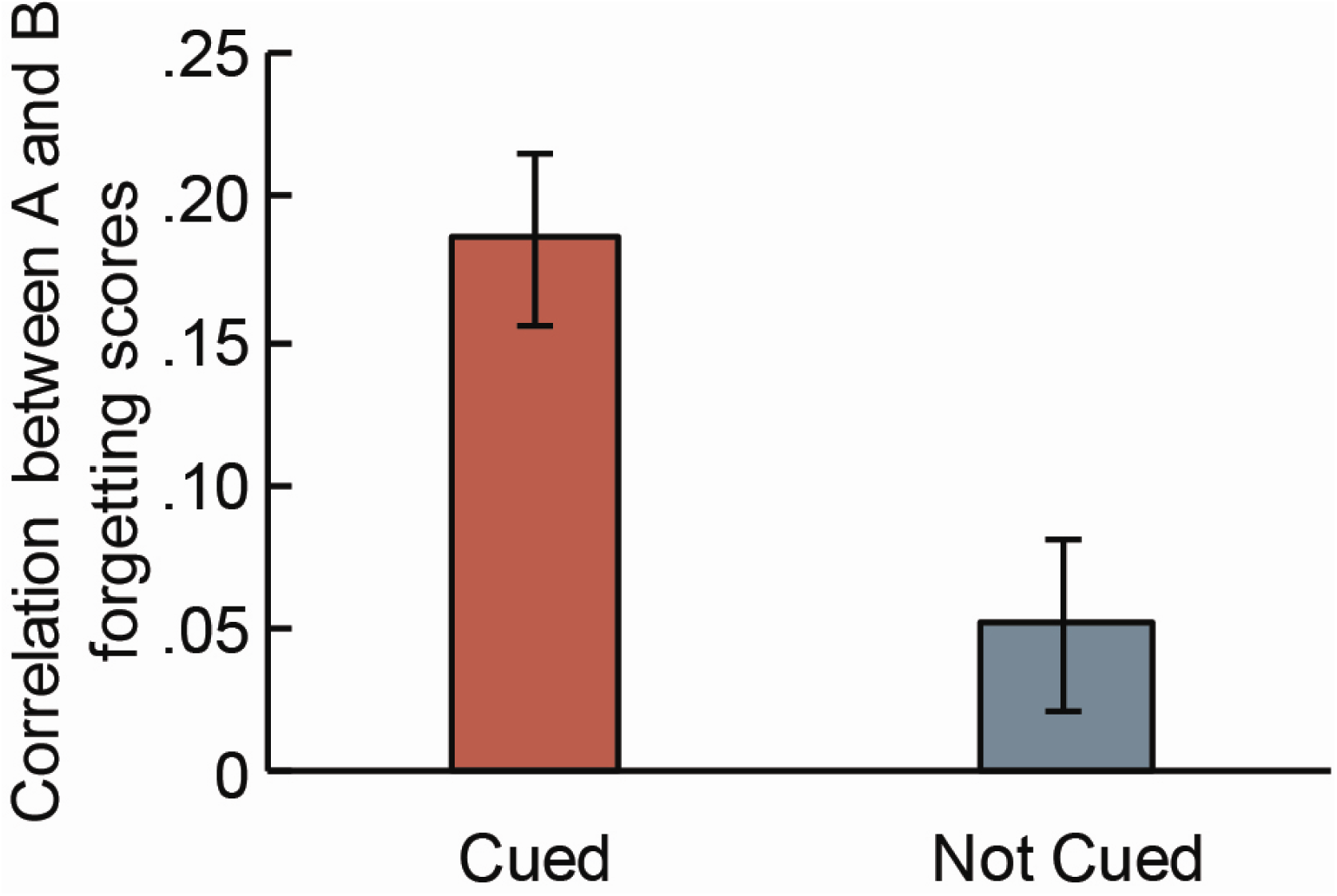
The correlation between the levels of forgetting of associated objects was marginally greater for pairs that were cued versus pairs that were not cued. Error bars represent ± 1 SEM (computed as in Figure 2).

Analysis of time in each sleep stage and correlations with behavior are presented in Table 3. There were no significant correlations between cuing advantage and time in each sleep stage. Additionally, we calculated the correlation between cuing advantage and delta power at the frontal electrode cluster (Fpz, Fp1, and Fp2) as well as the correlation between cuing advantage and slow spindle density at the frontal cluster and fast spindle density at the central-parietal cluster (Cz, Pz, C3, C4, P3, P4). None of the correlations with cuing advantage were significant (delta power: r = .02, *p* = .94; slow spindle density: r = −.19, *p* = .37; fast spindle density: r = .36, *p* = .11).

**Table 3.** Sleep physiology and correlations with behavioral data. * *p* < .05, † *p* < .10

|  |  | Cuing Advantage |  | Change in Error Correlation |  |
| --- | --- | --- | --- | --- | --- |
| Sleep Stage | Time in min<br>(mean ± SEM) | *r* | *p* | *r* | *p* |
| Wake | 24.1 ± 3.1 | -.38 | .09† | -.27 | .22 |
| Stage 1 | 4.6 ± .9 | -.26 | .24 | -.17 | .44 |
| Stage 2 | 26.1 ± 2.4 | .16 | .47 | -.05 | .81 |
| SWS | 22.8 ± 3.3 | .19 | .38 | .39 | .08† |
| REM | 4.1 ± 1.4 | -.24 | .28 | -.04 | .86 |
| Total Sleep | 57.6 ± 3.7 | .13 | .56 | .27 | .22 |

After removing data from an outlier participant who had excessive SWS (69 minutes, over 3 times the average, which was 22.8 minutes), an exploratory analysis showed a strong relationship between the change in correlation between A and B error and amount of SWS (r = .50, *p* < .03, Table 4). Additionally, there was a correlation between the primary two outcome measures and the average number of cues per pair received during sleep (Cuing Advantage: r = .46, *p* < .05, Figure 6; Change in Error Correlation: r = .44, *p* = .06). There was also a significant correlation between Change in Error Correlation due to cuing and delta power (r = .49, *p* < .05) as well as slow-oscillation density (r = .47, *p* < .05) at frontal electrode clusters (Fpz, Fp1, Fp2). Change in Error Correlation also showed a relationship with sigma power (r = .45, *p* < .05) and slow oscillation density (r = .45, *p* < .05) at central electrode clusters.

**Figure 6.**
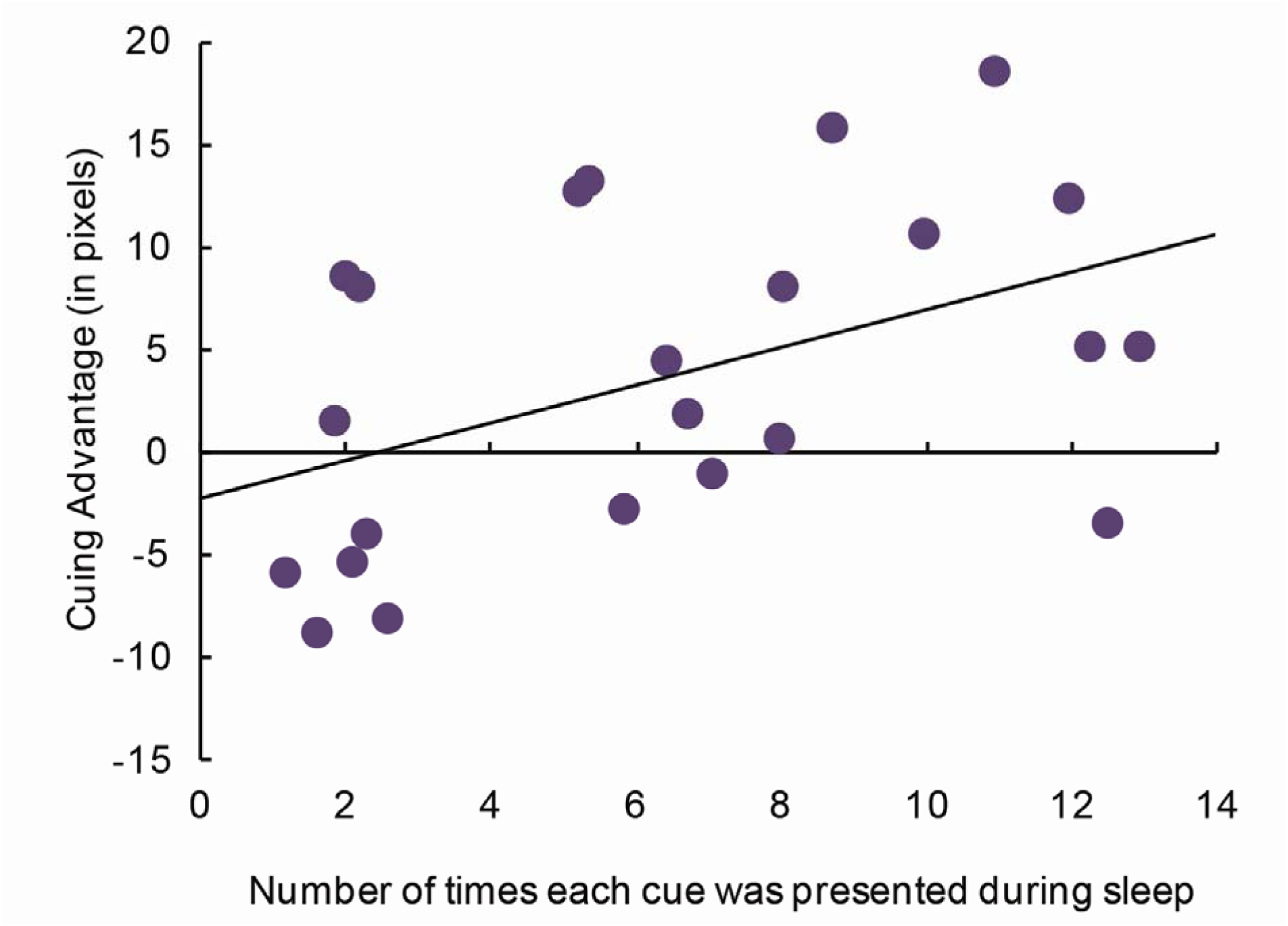
The more times each cue was presented during sleep, the less forgetting for cued compared to non-cued objects (r = .46, *p* < .05). This analysis excluded an outlier who had 69 minutes of SWS.

**Table 4.** Sleep physiology and correlations with behavioral data, excluding outlier. * *p* < .05, † *p* < .10

|  |  | Cuing Advantage |  | Change in Error Correlation |  |
| --- | --- | --- | --- | --- | --- |
| Sleep Stage | Time in min<br>(mean ± SEM) | *r* | *p* | *r* | *p* |
| Wake | 24.8 ± 3.1 | -.40 | .09† | -.28 | .21 |
| Stage 1 | 4.8 ± .9 | -.28 | .21 | -.17 | .44 |
| Stage 2 | 26.9 ± 2.4 | .15 | .51 | -.06 | .80 |
| SWS | 20.8 ± 3.3 | .28 | .20 | .50 | .03* |
| REM | 4.3 ± 1.4 | -.25 | .27 | -.04 | .86 |
| Total Sleep | 56.8 ± 3.7 | .14 | .52 | .28 | .22 |

## Discussion

The present experiment provided novel support for the conclusion that brain activity during sleep can impact subsequent memory ability. Results replicated findings from previous sleep studies (e.g., Rudoy et al., 2009; Creery et al., 2015) in showing that playing sound cues that had been associated with learning strengthened corresponding object-location memories in comparison to object locations that were not cued. Additionally, the findings showed that one cue can reactivate and strengthen more than one object-location association. Memory strengthening was measured in the form of reduced forgetting for the two independent object locations in each cued pair. Each pair included one object that was semantically related to the sound and one that was not. Thus, these results also showed that sounds can reactivate memories even for objects not semantically connected with the presented sound.

Another conclusion supported by the results is that cued objects were reactivated together as pairs, rather than individually and independently. When time passes, it is reasonable to expect memory accuracy to decline, and also for forgetting to vary between the members of a pair. This pattern of variance in within-pair forgetting was observed here in analyses of correlations between recall accuracy for the two objects of each pair, A and B. This correlation weakened after the nap only for non-cued pairs. Additionally, the correlation between forgetting for Object A and Object B was marginally higher for cued relative to non-cued pairs. In other words, the interrelationship between recall accuracy for A and B members of each pair was influenced by whether the corresponding sound was presented during sleep. Given that each cue was presented multiple times, it is possible that each presentation reactivated only one memory each time (either A or B), but concurrent reactivation of A and B is a more parsimonious explanation for the observed patterns of within-pair forgetting variance.

The relationships observed between behavioral measures and sleep physiology, although not universally strong, are consistent with the view that sleep reactivation played an active role in strengthening location memories for both objects. Increases in sleep physiology measures of slow oscillations (frontal delta power and slow oscillation density) and spindles (central sigma power) corresponded to increases in error correlations between paired objects. Both slow oscillations and sleep spindles have been implicated as part of the mechanism through which memory consolidation occurs during sleep (Diekelman & Born, 2010).

Whereas memories for object pairs in our study may have been reactivated concurrently, it would be interesting to determine whether there are cases in which TMR cues engage memory competition. For example, if one cue is associated with two separate memories, cuing during sleep may reactivate only one of the associated memories based on motivating factors such as believing one memory is more important to remember over the other. Such studies may help elucidate what is replayed and how competition operates (Antony, Cheng, Pacheco, Wang, Paller, & Norman, under review). Using TMR techniques may also provide insights into the content of reactivation, factors that influence reactivation, and a timeline for these processes (e.g., Cairney, Guttesen, El Marj, & Staresina, 2018).

Given the paired objects in this study, twice as much information was reactivated and strengthened as in previous studies using a similar task with roughly the same amount of sleep (Rudoy et al., 2009; Creery et al., 2015). In contrast to the effects on spatial recall, however, we were unable to examine whether TMR strengthened the learned associations between object pairs due to ceiling-level performance in recalling object names. Given that most experiences people have require multiple types of information to come together to form a rich and cohesive memory, additional studies are needed to understand how more complex memories may be strengthened during sleep. If it were possible to artificially reactivate different aspects of a memory using TMR, there could be ways to strengthen desirable features of a memory over unwanted ones.

The results of this study also raise the possibility of strengthening indirectly cued associations (i.e., second- or third-order associations), which could in turn promote relational binding. Previous work has shown that sleep helps promote item-integration (Dumay & Gaskell, 2007; Ellenbogen et al., 2007; Lau et al., 2010), so it is possible that integration may also be enhanced using TMR. The formation of new implicit associations during sleep has been shown using conditioning with aversive odors paired with the odor of cigarettes during sleep to reduce smoking behavior (Arzi et al., 2014). Results from Hauner and colleagues (2013) also suggest that associative learning can be altered during sleep. Participants underwent olfactory contextual fear conditioning, and during sleep were re-exposed with an odor previously associated with a conditioned stimulus and a mild electric shock. Re-exposure to the odor helped promote extinction for the conditioned fear response, possibly by forming a new association between the conditioned stimulus and the absence of a shock. Artificial memory formation during sleep is also possible in rodents; place cell activity was monitored and rewarding stimulation used to create new place-reward memories (de Lavilléon, Lacroix, Rondi-Reig, & Benchenane, 2013). After sleep, rodents exhibited goal-directed behavior indicative of memory for the new association. The extent to which new explicit memories can be produced during sleep in humans remains to be determined.

Our results show that auditory TMR can enhance memory in relation to more than a single item. Olfactory TMR has been consistently used to enhance memory for multiple items (e.g., Rasch et al., 2007), yet these studies commonly employ only one or two odors that may be associated with a learning context and not individual items. Here, each sound had a fairly specific association. Yet, with the present design we cannot determine whether each sound was associated independently with each of the two corresponding objects. Perhaps in some cases a sound instead reactivated only Object A, which in turn reactivated Object B, or a sound may have conjointly reactivated the association between the two objects that the participant created during learning. Future studies should further explore the hypothesis that a single sound can reactivate multiple, independent memories and reveal the boundary conditions for these associations, which may expose the properties of the neural infrastructure supporting memory consolidation during sleep. Pursuing these avenues of research may reveal the mechanisms of reactivation and open new paths towards the utilization of TMR for memory improvement.

## Acknowledgements

This work was supported by National Science Foundation grant number BCS-1461088. IMV was funded by an NSF Graduate Research Fellowship. ES was funded by the Human Frontier Science Program and the Zuckerman STEM Leadership Program.

